# Impaired pattern separation in Tg2576 mice is associated with hyperexcitable dentate gyrus caused by Kv4.1 downregulation

**DOI:** 10.1101/2021.02.17.431569

**Authors:** Kyung-Ran Kim, Yoonsub Kim, Hyeon-Ju Jeong, Jong-Sun Kang, Sang Hun Lee, Yujin Kim, Suk-Ho Lee, Won-Kyung Ho

## Abstract

Alzheimer’s disease (AD) is a progressive neurodegenerative disorder that causes memory loss. Most AD researches have focused on neurodegeneration mechanisms. Considering that neurodegenerative changes are not reversible, understanding early functional changes before neurodegeneration is critical to develop new strategies for early detection and treatment of AD. We found that Tg2576 mice exhibited impaired pattern separation at the early preclinical stage. Based on previous studies suggesting a critical role of dentate gyrus (DG) in pattern separation, we investigated functional changes in DG of Tg2576 mice. We found that granule cells in DG (DG-GCs) in Tg2576 mice showed increased action potential firing in response to long depolarizations and reduced 4-AP sensitive K^+^-currents compared to DG-GCs in wild-type (WT) mice. Among Kv4 family channels, Kv4.1 mRNA expression in DG was significantly lower in Tg2576 mice. We confirmed that Kv4.1 protein expression was reduced in Tg2576, and this reduction was restored by antioxidant treatment. Hyperexcitable DG and impaired pattern separation in Tg2576 mice were also recovered by antioxidant treatment. These results highlight the hyperexcitability of DG-GCs as a pathophysiologic mechanism underlying early cognitive deficits in AD and Kv4.1 as a new target for AD pathogenesis in relation to increased oxidative stress.

## Introduction

Alzheimer’s disease (AD) is a progressive neurodegenerative disorder and is the most common cause of dementia. Considerable evidence has indicated that AD patients exhibit cognitive decline several years before clinical diagnosis (1-3). In particular, assessment of episodic memory is the most effective method for identifying at-risk individuals (3, 4). Synaptic loss associated with a decline in basal synaptic transmission and a deficit in induction of long-term potentiation (LTP) have been identified as potential underlying mechanisms for AD-associated cognitive declined (5-7). Synaptic loss and deficits were shown to be mediated by amyloid β (Aβ) and Tau oligomers, or epigenetic alterations (7). On the other hand, AD pathogenesis also exhibits a profound increase in hyperactive neurons (8, 9). Aberrant network activity associated with increased seizure are also considered as important factors causing cognitive decline (10). It is not yet fully understood how Aβ and Tau are involved in both synaptic depression and neuronal hyperexcitability or how they are related to aberrant network activity and cognitive decline.

Synaptic dysfunction is widely thought to be one of the earliest key pathogenic events in AD before frank neurodegeneration, and the hippocampal network is particularly vulnerable in AD (11-14). The hippocampus is divided into three main fields, the dentate gyrus (DG) and areas CA3 and CA1, and each field displays distinctive anatomical, molecular, and biophysical properties (15, 16). Although all three hippocampal pathways have been associated with learning and memory, MF-CA3 projection specifically has been implicated in cognitive function, including novelty detection, pattern completion and pattern separation (17, 18). Indeed, structural and functional MRI analysis of AD patients revealed disruption of the mossy fiber (MF)-CA3 pathway in patients with mild AD or mild cognitive impairment (19, 20). Synaptic activity in the DG/CA3 network was proposed as an early target of amyloid pathology that leads to impaired pattern separation and episodic memory loss (21). Consistently, impaired synaptic plasticity in MF-CA3 synapses at an early stage in AD model mice was reported (22, 23). However, it remains to be elucidated how alteration of intrinsic excitability contributes to impaired DG/CA3 network underlying AD-associated cognitive deficit

Sparseness of DG activity is a prerequisite for orthogonal representation of information which is critical for pattern separation function (24, 25). This hypothesis was supported by an experiment showing that hyperexcitability of GCs in DG induced by selective degeneration of hilar mossy cells led to impaired pattern separation (26). We have recently reported that low excitability of mature GCs is attributable to a high level of Kv4.1 expression, and knock-down of Kv4.1 in DG causes hyperexcitability and impaired pattern separation (27). In the present study, we investigated the possible role of Kv4.1 in impaired pattern separation in the early preclinical stage of AD, and found that GCs in Tg2576 mice showed hyperexcitability and reduced Kv4.1. Furthermore, hyperexcitability and reduced Kv4.1 in Tg2576 GCs and impaired pattern separation in Tg2576 mice were reversed by antioxidant treatment. These results highlight the hyperexcitability of GCs in DG as a pathophysiologic mechanism underlying early cognitive deficits in AD, and Kv4.1 as a new target for AD pathogenesis in relation to increased oxidative stress.

## Results

### Impaired pattern separation in Tg2576 mice

To investigate whether AD model mice in an early stage before expressing amyloid plaque deposition and severe memory loss have impairment in pattern separation, we used 3 ∼ 4 months-old Tg2576 mice and performed contextual fear discrimination tests (28) (Fig. 1A). A similar pair of contexts (A and B) had identical metal grid floors, while context B had a unique odor (1 % acetic acid) with dimmer lighting (50 % of A) in the sloped floor (15° angle). As shown in Figure 1B, mice are taught to distinguish between the similar contexts over several days. We observed that both WT and Tg2576 groups showed comparable freezing levels during 5 min test in context A (Fig. 1Ba). Moreover, freezing levels were not different between context A and B in both genotypes (Fig. 1Bb, WT, Tg2576, two-way ANOVA, genotype: F_(1,30)_ = 0.24, p = 0.62, context: F_(1,30)_ = 0.15, p = 0.70, genotype × context: F_(1,30)_ = 0.39, p = 0.54). The mice were subsequently trained to disambiguate these contexts by visiting both contexts daily for 8 days with a 2-hr-interval between the contexts (from day 6 to day 13), always receiving a foot shock 180 s after being placed in context A but not in context B. A daily discrimination ratio was calculated by freezing during the 180 s in context A over total freezing during the two visits (A and B). On day 6, both groups could not distinguish the differences between the contexts (Fig. 1Cb, genotype: F_(1,24)_ = 3.67, p = 0.06, context: F_(1,24)_= 0.73, p = 0.40, genotype × context: F_(1,24)_ = 0.31, p = 0.58); thus, the discrimination ratio was approximately 0.5. Over subsequent training days, WT mice began to discriminate context B from context A with an increase in discrimination ratio. However, Tg2576 mice exhibited significant deficits in the acquisition of discrimination ability (Fig. 1Ca), and showed increased freezing in the shock-free context (Fig. 1Cb, t-test, Context B, p < 0.0001, Two-way ANOVA, genotype: F_(1,25)_= 0.25, p = 0.62, context: F_(1,25)_= 19.19, p < 0.0001, genotype × context: F_(1,25)_ = 34.63, p < 0.000004). To rule out the possibility that general memory acquisition or storage is impaired in Tg2676 mice, the context specificity of the conditioning was examined using context A and context C that is distinct from context A. In this configuration, context C evoked a significantly lower level of freezing than context A in both genotypes (Fig. 1D, genotype: F_(1,14)_= 0.16, p = 0.70, context: F_(1,14)_= 43.78, p < 0.000012, genotype × context: F_(1,11)_ = 0.06, p = 0.80). These results indicate that 3 ∼ 4-month-old Tg2576 mice exhibit specific impairment in pattern separation between similar contexts without deficits in memory acquisition or learning.

**Figure 1.**
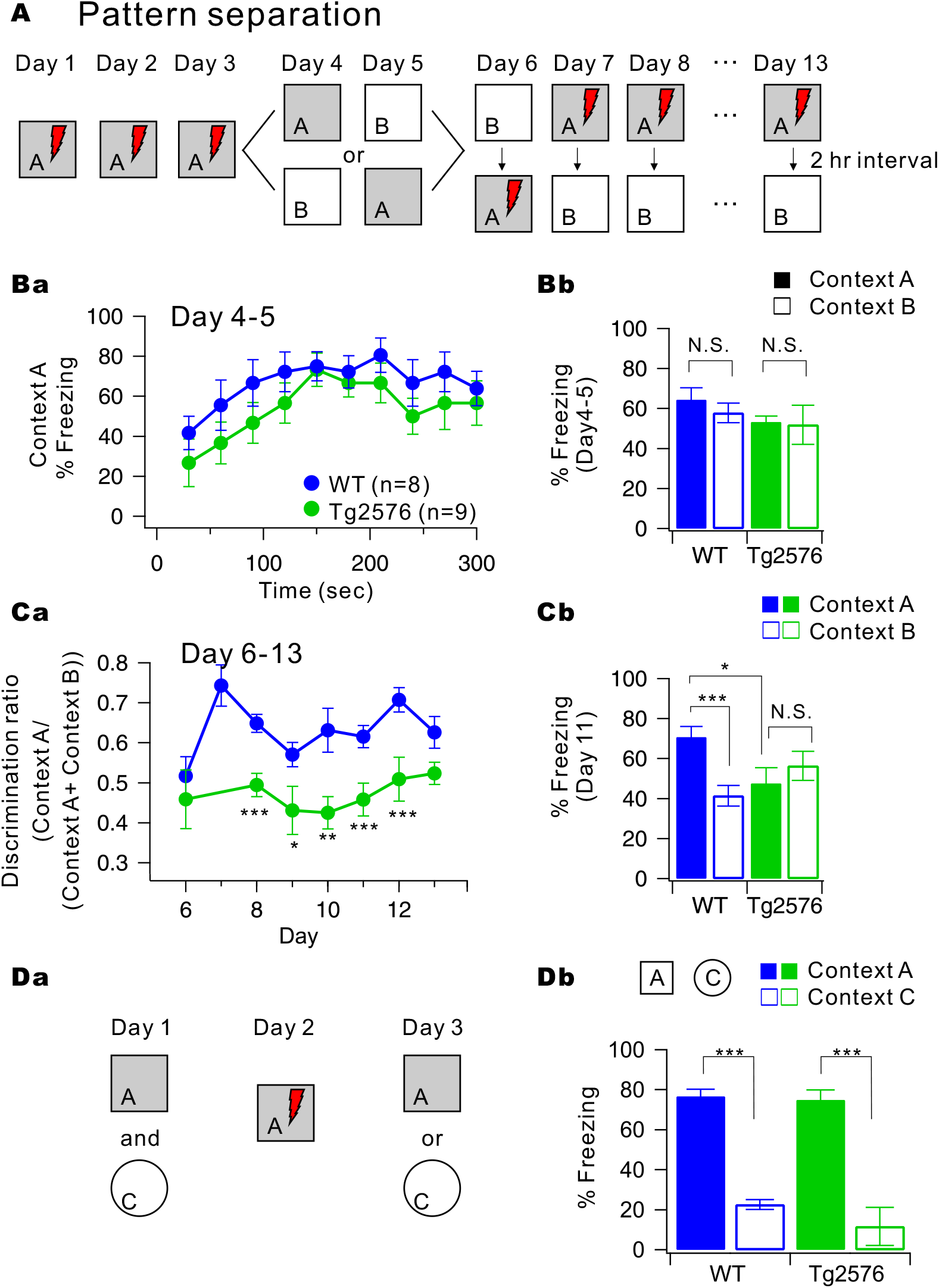
Impaired pattern separation in Tg2576 mice. ***A***, Experimental procedure for pattern separation in 10-week-old WT and Tg2576 mice. ***Ba***, On day 4 to 5, the kinetics of freezing across the 5 min test in context A. ***Bb***, Percentage of freezing in context A (filled bar) and context B (open bar) during day 4 to 5. WT (blue, *n* = 8) and Tg2576 (green, *n* = 9) mice displayed equal amounts of freezing in both contexts A and B. ***Ca***, On day 6 to 13, time course of the discrimination ratio in WT (blue, *n =* 8) and Tg2576 (green, *n* = 7) mice. Statistical significance for each day was tested using ANOVA (\**P* < 0.05; \*\**P* < 0.01; *** *P* < 0.001). ***Cb***, Percentage of freezing in context A (filled bar) and context B (open bar) for WT (blue, *n* = 8) and Tg2576 (green, *n* = 7) mice on day 11. Freezing levels were compared between the two contexts for each group (N.S., no statistical significance; \**P* < 0.05). ***Da***, Experimental procedure for one-trial contextual fear conditioning between WT (*n* = 8) and Tg2576 (*n* = 7) mice. ***Db***, The percentage of freezing in context A (filled bar) and context C (open bar, distinct object) for WT (*n* = 8) and Tg2576 (*n* = 7) mice.

### Hyperexcitability and reduced K^+^ currents in Tg2576 GCs in DG

To investigate the possibility that the abnormal excitability of granule cells (GCs) in dentate gyrus (DG) is involved in the impairments of pattern separation in Tg2576 mice, we examined intrinsic properties of GCs. To avoid the neural progenitor cells and newborn immature neurons, we identified mature DG granule neurons in the hippocampus according to their morphological and electrophysiological criteria: low resting membrane potential (RMP) and input resistance (R_in_; <200 MΩ) located in the outer molecular layer (27, 29). We found that firing frequency evoked by depolarizing pulses of 1 sec duration was significantly higher in the Tg2576 GCs (green, Fig. 2A) compared to the WT GCs (blue, Fig. 2A). The relationship of action potential (AP) frequency (F) versus injected currents (I) showed an upward shift in DG neurons of Tg2576 mice relative controls, indicating increased neuronal excitability (Fig. 2Ab). In agreement with these findings, the AP onset time during AP trains was also significantly shorter in Tg2576 GCs (Fig. 2B; 34.7 ± 1.3 ms, *n* = 13, for WT and 24.6 ± 1.6 ms, *n* = 12, for Tg2576 at 300 pA current injection, p=0.000045). However, we did not find significant differences between Tg2576 and WT mice in passive electrical properties, such as RMP, R_in_ and threshold current for the AP generation (Rheobase) (Fig. 2C; RMP: p=0.,11641, R_in_: p=0.8121, Rheobase: p=0.46504). However, unlike DG GCs, the excitability of CA1 pyramidal neurons was not different between Tg2576 and WT mice (Fig. 2D), suggesting the GC-specific hyperexcitability in Tg2576 mice. These results suggest that the enhanced excitability of GCs in Tg2576 mice is unlikely to be attributable to the alteration of Na^+^ channels or ion channels in determining the subthreshold membrane excitability. We therefore hypothesized that downregulation of Kv4.1 channels that play key roles in low excitability of mature GCs (27) may cause the hyperexcitability of Tg2576 GCs..

**Figure 2.**
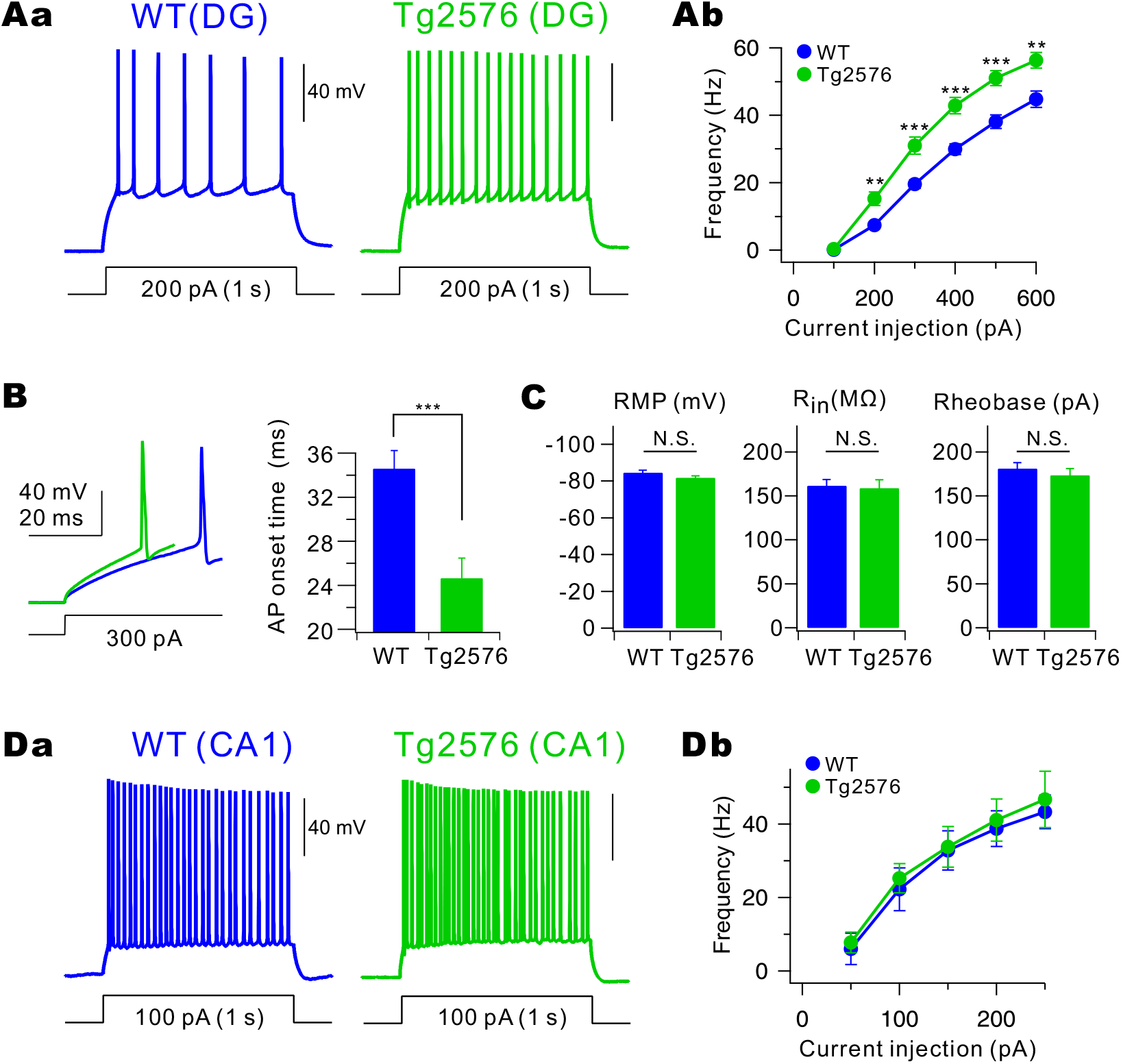
Electrophysiological analysis for the excitability of WT and Tg2576 mice. ***Aa***, Representative traces in the current-clamp recording from the WT (blue) and Tg2576 (green) dentate GCs in response to prolonged (1 s) depolarizing current injection (200 pA). ***Ab***, Number of spike evoked during 1 s current injections with varying amplitudes (from 100 pA to 600 pA, 100 pA increment). At all amplitudes, the number of spikes is significantly higher in Tg2576 GCs (green closed circle, *n* = 14) compared to WT GCs (blue closed circle, *n* = 15). Mean ± S.E.M. \*\**P* < 0.01; *** *P* < 0.001 ***B***, Expanded view of overlaid 1st APs evoked by 300 pA current injection to show the shortening of AP onset in Tg2576 GCs compared to WT GCs. Mean ± S.E.M. *** *P* < 0.001 ***C***, The mean values of the resting membrane potential (WT: −84.5 ± 1.3 mV, Tg2576: −81.8 ± 1.0 mV), input resistance (WT: 161.6 ± 6.9 MΩ, Tg2576: 158.8 ± 9.5 MΩ), and rheobase (WT: 181.1 ± 6.7 pA, Tg2576: 173.5 ± 7.7) from the WT (*n* = 15) and Tg2576 GCs (*n* = 14). Mean ± S.E.M. N.S., not significantly different. ***Da***, Representative traces of current-clamp recordings from CA1 of WT (blue) and Tg2576 mice (green). A train of APs were evoked by 1 s depolarizing current pulse injection (100 pA). ***Db***, The F-I curve for CA1 from WT (*n* = 4) and Tg2576 (*n* = 4) mice.

To examine this hypothesis, we first recorded K^+^ currents in the voltage clamp mode by applying depolarizing voltage steps (ranging from −60 mV to +30 mV with 10 mV increment and 1 sec duration) at a holding potential of −70 mV. TTX, Cd^2+^/Ni^2+^, bicuculline, and CNQX were added to the external solution to block Na^+^ channels, Ca^2+^ channels, GABA_A_ receptors, and AMPA/kainate receptors, respectively. We found that total outward K^+^ currents in Tg2576 GCs are similar to that in WT GCs (Fig. 3A), while K^+^ currents sensitive to 5 mM 4-AP (I_4-AP_) are substantially less in Tg2576 GCs relative to WT GCs (Fig. 3B). The difference was found both in the peak current (I_peak_) (at +30 mV; WT: 2.234 ± 0.203 nA, *n* = 9, Tg2576; 1.624 ± 0.197 nA, *n* = *7*, p = 0.0477) and in the steady state current (I_ss_) (at +30 mV; WT: 0.505 ± 0.042 nA, *n* = 9, Tg2576: 0.161 ± 0.037 nA, *n* = 7, p = 0.000197). Notably, reduction of I_ss_ was greater than I_peak_ in Tg2576, suggesting that reduced I_4-AP_ in Tg2576 GCs is attributable to downregulation of K^+^ channels that inactivate slowly. We further analyzed the difference in I_4-AP_ between WT and Tg2576 GCs (Fig. 3C). To further analyze inactivation of I_4-AP_, we fitted inactivation phases of I_4-AP_ with a two-exponential decay function, I(t) = A_fast_ exp(-t/τ_fast_) + A_slow_ exp(-t/τ_slow_). The results showed that the amplitude of the slow component (A_slow_) was selectively reduced in the Tg2576 GCs compared to WT GCs, whereas the other parameters (A_fast_, τ_fast,_ and τ_slow_) were not altered (Fig. 3C). Considering that the slowly inactivating component of I_4-AP_ in GCs was mainly attributable to Kv4.1 current (27), these results suggest that reduced I_4-AP_ in Tg2576 GCs is attributable to the decrease in Kv4.1 current.

**Figure 3.**
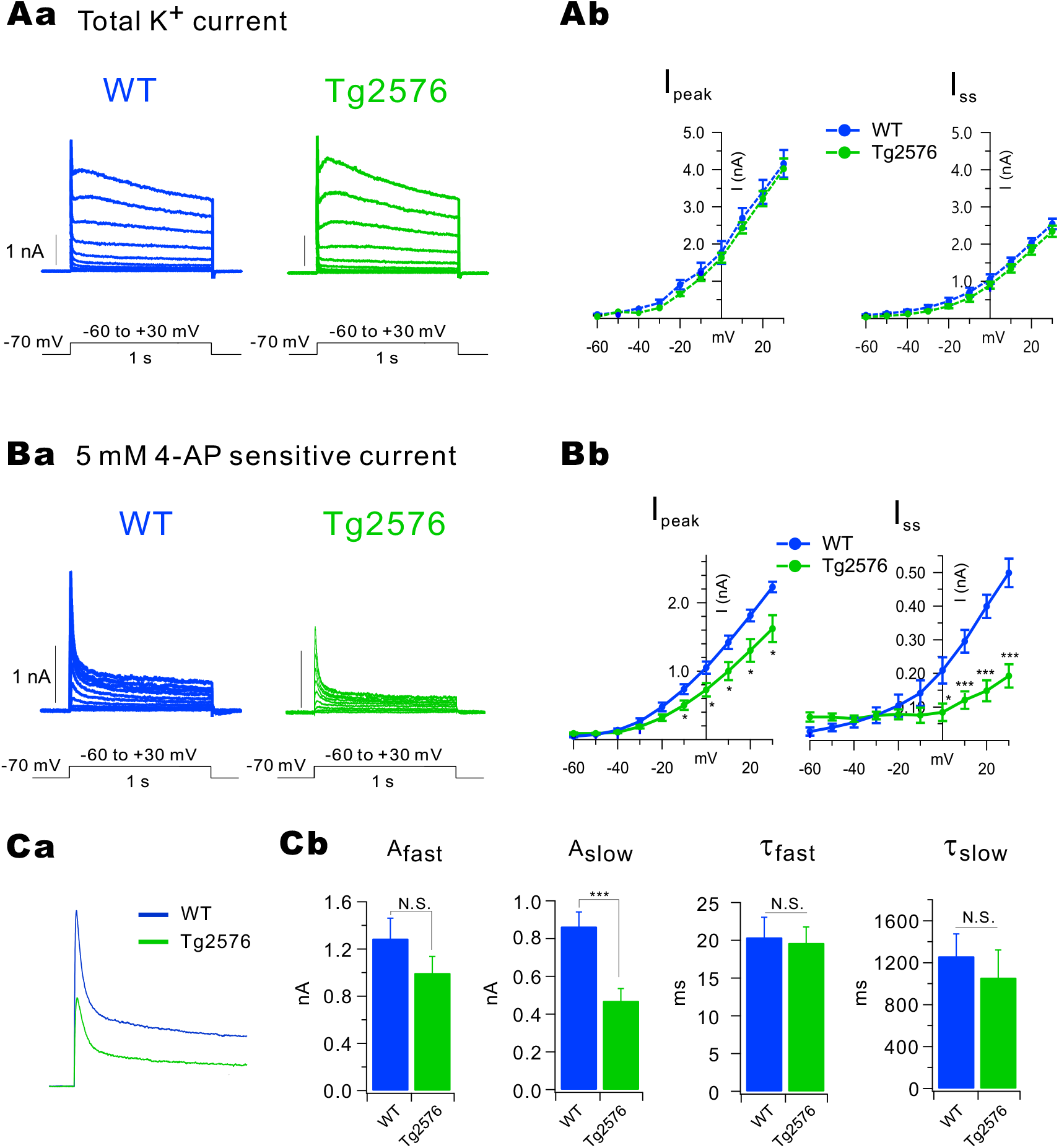
Voltage-dependent K^+^ currents in the WT and Tg2576. ***Aa***, Superimposed current traces evoked by various steps of depolarization (−60 mV to +30 mV in 10 mV step) for 1 s from the holding potential of −70 mV in the WT (blue) and Tg2576 (green) dentate GCs. ***Ab***, Amplitudes of peak currents (I_peak_) and steady state currents (I_ss_) measured at the end of depolarization are plotted as a function of the given potential (V). WT (closed blue circle, *n* = 9) and Tg2576 (closed green circle, *n* = 7). ***Ba***, The difference in currents before and after applying 5 mM 4-AP (I_4-AP_) in the WT (blue) and Tg2576 (green) dentate GCs. ***Bb***, I_peak_ and I_ss_ for I_4-AP_ are plotted as a function of the given potential (V). WT (closed blue circle, *n* = 9) and Tg2576 (closed green circle, *n* = 7). ***Ca***, I_4-AP_ traces obtained at +30 mV are averaged (WT, blue, *n* = 9; Tg2576, green, *n* = 7) and superimposed. ***Cb***, I_4-AP_ recorded at +30 mV were fitted to double-exponential functions. Amplitude and time constant for the fast component (A_fast_, τ_fast_) and the slow component (A_slow_, τ_slow_). Mean ± S.E.M., **, *P* < 0.01, ***, *P* < 0.001. N.S., not significantly different, *P* > 0.05.

### Kv4.1 downregulation is responsible for hyperexcitability of Tg2576 GCs

To investigate the molecular identity of the decreased I_4-AP_ in Tg2576 GCs, we examined whether mRNA expression levels of Kv4 family channels are altered using quantitative Real-Time PCR (qRT-PCR) techniques. We found that the Kv4.1 mRNA expression level in DG was significantly lower in Tg2576, while Kv4.2 and Kv4.3 mRNA expressions levels were not altered (Fig. 4A; Kv4.1: p=0.000058, Kv4.2: p=1, Kv4.3: p=0.62857, Mann-Whitney test). These results suggest that Kv4.1 expression was selectively downregulated in Tg2576 without compensation by the other Kv4 channel subunits. In agreement with these findings, Kv4.1 protein level in DG was also significantly lower in Tg2576 compared to WT (Fig. 4B). To examine the functionality of reduced Kv4.1 expression in Tg2576, we examined the effects of Kv4.1 antibody perfusion via a whole-cell patch pipette on firing frequency. In WT GCs, action potential firing evoked by depolarizing pulses of 1 sec duration was increased significantly by Kv4.1 antibody (from 7.6 ± 0.9 to 14.2 ± 3.2, *n* = 5, p = 0.00718, Fig. 4C), which is consistent with our previous results in GCs of C57BL/6 control mice (27). These results confirmed that Kv4.1 contributes to lowering firing frequency in GCs in WT as in C57BL/6 mice. In contrast, firing frequency was not significantly affected by Kv4.1 antibody in Tg2576 GCs (15.2 ± 1.9 *vs* 17.3 ± 2.9, p = 0.64891, Fig. 4C). It was also noted that firing frequency of WT GCs with Kv4.1 antibody was comparable with that of Tg2576 GCs (p = 0.79502, Fig. 4C). These results support the idea that Kv4.1 is responsible for limiting AP firing in WT GCs, and that reduced Kv4.1 in Tg2576 GCs is a key mechanism for enhanced excitability in Tg2576 GCs.

**Figure 4.**
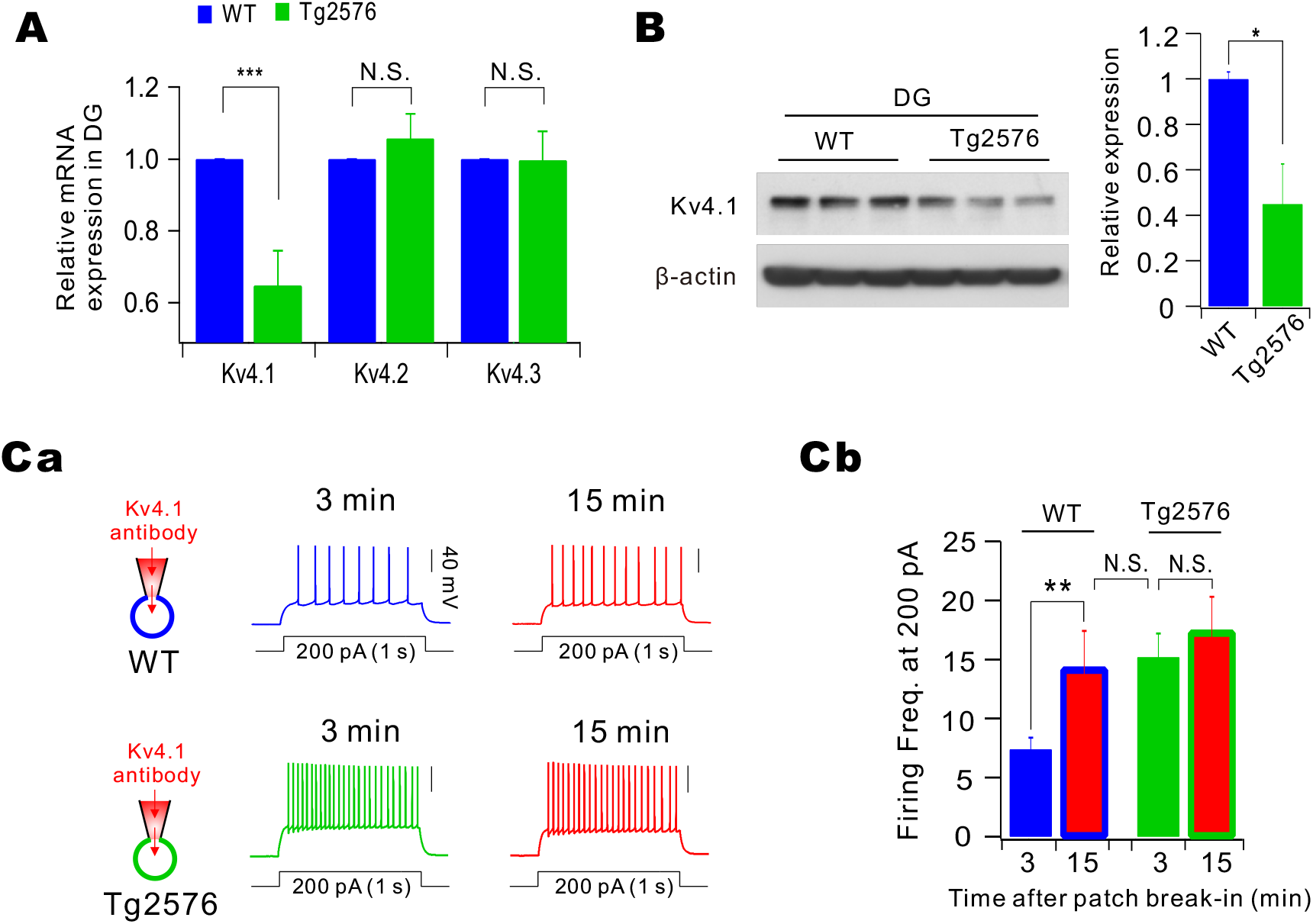
Downregulation of Kv4.1 expression in Tg2576 causes hyperexcitability of Tg2576 GCs. ***A***, mRNA expression levels for Kv4 family channels in the WT (blue) and Tg2576 (green) DG detected by quantitative RT-PCR. ***B***, Western blot analysis for Kv4.1 in DG of 8-week-old mice. For a loading control, b-actin was used. Right, Kv4.1expression in Tg2576 was significantly smaller than WT mice. *, *P* < 0.05. ***Ca***, Example traces of APs evoked by 200 pA current injection immediate (3 min, blue for WT and green for Tg2576) and 15 min (red) after brake-in of patch pipettes containing Kv4.1 antibody. ***Cb***, Firing frequency at 200 pA is compared in each conditions.

### Hyperexcitability of Tg2576 GCs and impaired pattern separation in Tg2576 mice are restored by antioxidant treatment

We previously reported that mitochondrial reactive oxygen species (ROS) production is increased in Tg2576 DG at the age of 1- to 2-months, causing depolarization of mitochondrial membrane potential and impairment of Ca^2+^ uptake int mitochondria. Furthermore, this impairment is restored by treatment of antioxidant, Trolox (30). We addressed the question whether ROS overproduction underlies the reduction of Kv4.1 expression and hyperexcitability in Tg2576 GCs. To this end, we first examined the effects of Trolox on the hyperexcitability phenotype in the Tg2576 GCs. Preincubation of Tg2576 GCs in the aCSF containing Trolox (500 μM) for 1 hr did not affect the relationship of AP frequency and injected currents (orange, Fig. 5A), while injection of Trolox (20 mg/Kg) into Tg2576 mice intraperitoneally (i.p.) once a day for one week induced a downward shift in the relationship of AP frequency and injected currents (purple, Fig. 5A). AP frequency in Tg2576 GCs following Trolox treatment for one week is similar to that in WT GCs (dashed blue line, Fig. 5Ab). Consistently, Western blot analysis for Kv4.1 proteins showed a significant increase in Tg2576 mice after one week treatment of treated with Trolox (Fig. 5B). These results suggest that restoration of excitability and Kv4.1 expression by Trolox treatment is not a rapid process. We finally examined whether Trolox treatment restored impaired pattern separation in Tg2576 mice. Contextual fear discrimination tests described in Fig. 1 were performed in Tg2576 mice treated with Trolox for one week. Freezing levels during 5 min test at day 4-5 showed comparable freezing levels with those observed in Tg2576 mice (Fig. 5C), while deficits in contextual fear discrimination observed in Tg2576 mice (green, Fig. 5D) was restored by Trolox treatment (purple, Figs. 5D). These results support the idea that impaired pattern separation in the early preclinical stage of AD is attributable to the hyperexcitability of GCs mediated by ROS-induced downregulation of Kv4.1.

**Figure 5.**
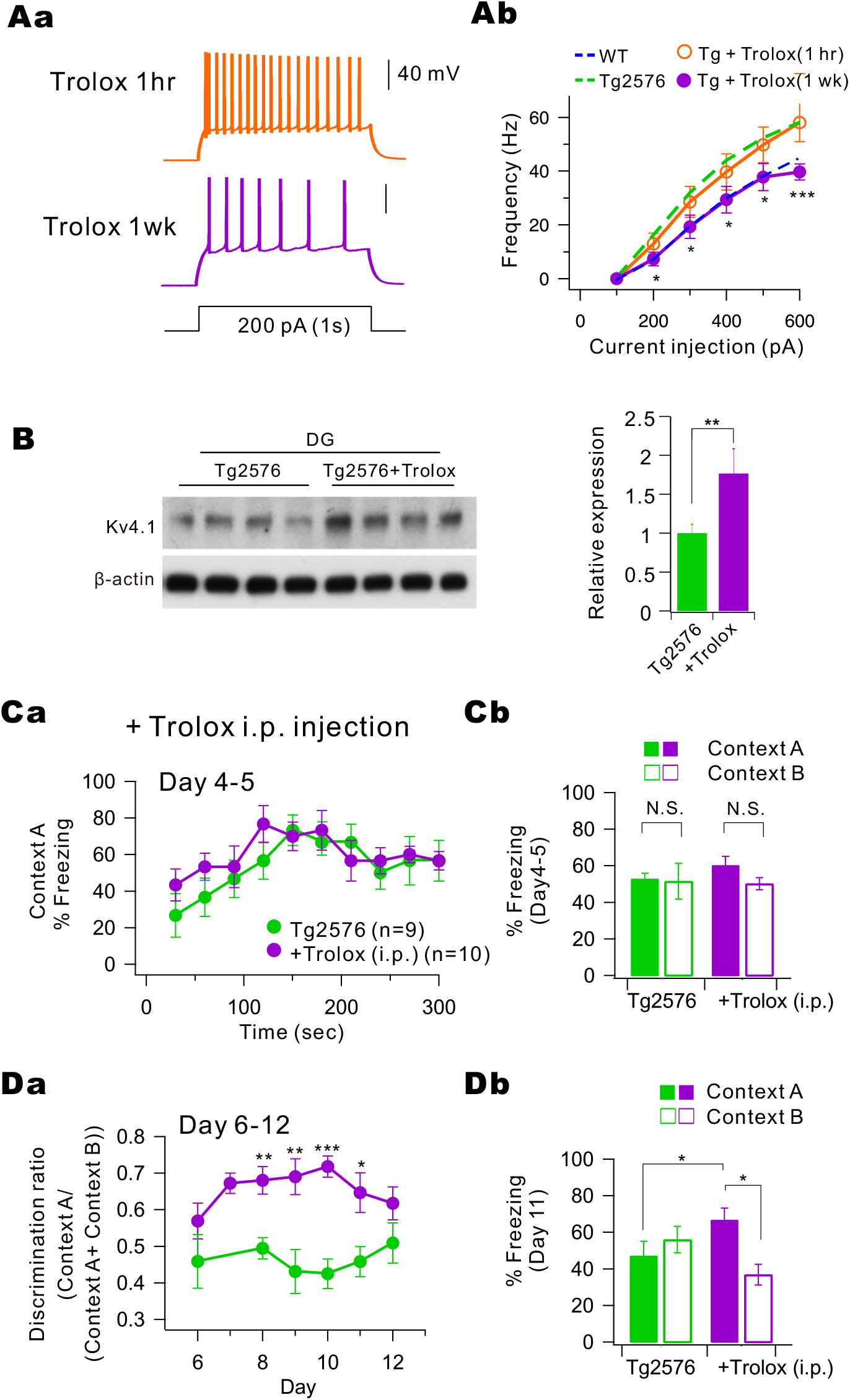
Effect of pre-treatment with Trolox in the Tg2576 mice. ***Aa***, Representative trace in the current-clamp recording of pre-treatment with Trolox for one hour (orange) and one week (purple) in Tg2576 in response to depolarizing current injection (200 pA with 1 s duration). ***Ab***, F-I curve (from 100 pA to 600 pA, 100 pA increment) obtained from GCs of Tg2576 mice treated with Trolox for one week (Tg+Trolox, purple, *n* = 8) was compared with the F-I curve obtained from Tg2576 GCs (green dashed line, *n* = 14) shown in Fig. 2Ab. *, *P* < 0.05, ***, *P* < 0.001. ***B***, Western blot analysis for Kv4.1 in DG of 8-week-old Tg2576 mice and those treated with Trolox (20 mg/kg) for one week. For a loading control, β-actin was used. Right, Kv4.1expression in Tg2576 was significantly increased after Trolox treatment. **, *P* < 0.01. ***Ca***, On day 4 to 5, the kinetics of freezing across the 5 min test in context A obtained from Tg2576 mice treated with Trolox for one week (Tg+Trolox, purple, *n* = 10) was compared with the that obtained from control Tg2576 GCs (green, *n* = 7) shown in Fig. 1Ba. ***Cb***, Percentage of freezing in context A (filled bar) and context B (open bar) during day 4 to 5. Tg2567 (green, *n* = 7) and Tg+Trolox (purple, *n* = 10) mice displayed equal amounts of freezing in both contexts A and B. ***Da***, On day 6 to 12, time course of the discrimination ratio in Tg+Trolox mice (purple, *n* = 10) was compared with that obtained from control Tg2576 mice shown in Fig. 1Ca (green, *n* = 7) mice. Statistical significance for each day was tested using unpaired *t*-test (\**P* < 0.05; \*\**P* < 0.01; *** *P* < 0.001). ***Db***, Percentage of freezing in context A (filled bar) and context B (open bar) for Tg2576 (green, *n* = 7) and Tg+Trolox (purple, *n* = 10) mice on day 11. Freezing levels were compared between the two contexts for each group. Tg2567 (green, *n* = 7) mice displayed equal amounts of freezing in both contexts A and B, while and Tg+Trolox (purple, *n* = 10) mice displayed more freezing in context A compared to context B.

## Discussion

AD is characterized as a neurodegenerative disease, although functional deficits occurred prior to neurodegeneration. We reported in our previous study that impaired post-tetanic potentiation was observed at the preclinical stage of AD (22). It was demonstrated that mitochondrial dysfunctions such as increased ROS production, partial depolarization of mitochondrial potential, and impaired Ca^2+^ uptake occurred in dentate GCs of 1∼2-month-old Tg2576 mice, leading to the deficits in mitochondria-dependent short-term plasticity at the mossy fiber-CA3 synapses. The observation that antioxidant treatment can restore the reduced short-term plasticity and impaired mitochondrial Ca^2+^ uptake in Tg2576 mice suggests that the increased ROS production in mitochondria initiates the other dysfunctions. In the present study, we revealed that hyperexcitability of GCs caused by Kv4.1 downregulation occurs in the early stage of AD. Furthermore, we showed that impaired pattern separation, possibly an early symptom of cognitive deficit, is linked to hyperexcitability of GCs and that both can be restored by antioxidant treatment. Our study suggests that ion channel remodeling induced by oxidative stress may underlie cognitive deficit in early AD pathogenesis.

Elevated excitability in the hippocampus has emerged as a key contributor to cognitive impairment in AD, though some studies reported reduced intrinsic excitability associated with AD (31, 32). Hyperexcitability including increased firing rate and an elevated number of activated neurons associated with AD was reported in CA1 (33-35) and CA3 pyramidal neurons (36, 37). Tight coupling between excitability and cognitive phenotype was also recognized in aging (38). Reducing hyperexcitability with the antiepileptic drug levetiracetam restores cognitive deficits in AD as well as in mild cognitive impairments (39-41), indicating the importance of hyperexcitability in AD pathogenesis. However, ion channel mechanisms underlying hyperexcitability were not identified in most studies. Kv4.2 was suggested to be involved in altered excitability in pathologic condition, yet the role is not conclusive. Kv4.2 current was reduced by intracellular accumulation of Aβ42 (35) or Tau (42), while an aging-related increase in A-type potassium currents was attributed to increased firing frequency by facilitating repolarization (37). In the present study, we not only demonstrated hyperexcitability of GCs in AD model mice, but also identified Kv4.1 downregulation in GCs as underlying mechanism of hyperexcitability. Causal relationship between decreased Kv4.1 expression, hyperexcitability of GCs, and impaired pattern separation in Tg2576 mice is strongly supported by our previous study (27).

Among many complex factors underlying AD pathogenesis, network abnormality caused by neuronal hyperactivity has recently emerged as a potential mechanism of cognitive dysfunction (10, 43-45) and epileptic seizure (44, 46-48). A causal link between Aβ accumulation and network abnormalities and epileptogenesis was proposed (44, 48). Functional magnetic resonance imaging (fMRI) of humans with early symptomatic AD has demonstrated excessive neuronal activities in the hippocampus and neocortex, where Aβ accumulates in abundance (9). Since neuronal hyperactivity increases extracellular Aβ levels (49), it may initiate a vicious cycle that facilitates progressive pathology and memory impairment (50). The molecular basis of this hyperexcitability has been examined, and Aβ-induced hyperexcitability of layer 2/3 (L2/3) pyramidal neurons (43), the dysfunction of parvalbumin-positive interneurons in L2/3 parietal cortex (51), and impaired glutamate reuptake by neurons and astrocytes (48) have been proposed so far. Considering that DG functions as a “gate” controlling information flow from the entorhinal cortex into the rest of the hippocampus, and hyperexcitability of dentate GCs has long been known as one of epileptogenic mechanisms underlying temporal lobe epilepsy (52, 53), hyperexcitability of GCs caused by downregulation of Kv4.1 may contribute to network abnormalities associated with increased seizure as well as cognitive impairment associated in AD. Importance of DG in AD pathogenesis has been recently highlighted in studies showing restoration of memory in AD mice by optogenetic activation of DG engram cells (54, 55). DG manipulation may be a potential target for treating cognitive deficit and memory loss in AD.

Kv4.1 belongs to the Kv4 gene subfamily together with Kv4.2 and Kv4.3 (56-58). Based on electrophysiological and pharmacological properties, Kv4 channels are known to mediate A-type K^+^ currents (I_A_), which are characterized by their rapid inactivation kinetics and their sensitivity to 4-aminopyridine (4-AP) blockade. However, a recent study showed that Kv4.1 in the hippocampus has unique electrophysiological characteristics and expression patterns distinctive from those of Kv4.2 (27). Kv4.2 was shown to mediate I_A_ in CA1 pyramidal neurons, regulating AP repolarization phase (59-61), while Kv4.1 was weakly expressed and had no role in CA1 pyramidal neurons (27). Interesingly, Kv4.1 expression was significant in DG, and Kv4.1-mediated currents detected only in low-frequency firing mature GCs displayed slow inactivation kinetics (27). Inhibition of Kv4.1 currents by shRNA or antibody induced a significant increase in firing frequency (27), suggesting that Kv4.1 played a key role in lowering firing frequency in mature GCs. Furthermore, specific knockdown of Kv4.1 in the DG region selectively impaired conditional freezing between similar contexts, suggesting that low frequency firing of mature GCs is crucial for pattern separation (27). The present study highlights Kv4.1 as a target ion channel that causes hyperexcitability of DG in AD, leading to impaired pattern separation.

Our previous study suggested that dentate GCs may be the earliest target for AD-associated oxidative stress (22). The present study implies that overproduction of ROS is responsible for the reduction of Kv4.1 expression in the young Tg2576 mice. The molecular mechanism how oxidative stress involved in AD pathology decreases Kv4.1 expression remains to be investigated. ROS-dependent regulation of channel expression is little understood in neurons, whereas a few studies in non-neuronal cells showed that ROS exerts diverse effects on expression and activity of Na^+^, K^+^ and Ca^2+^ channels (62-64). Mitochondria are not only the major source of ROS but also the primary target of ROS, which induces cellular dysfunction. Overproduction of ROS in Tg2576 mice is correlated with mitochondrial Aβ accumulation, probably leading to mitochondrial depolarization and the reduction in respiratory enzyme activities and oxygen consumption (22, 65, 66). The treatment of Trolox on GCs of Tg2576 mice restored the mitochondrial function in parallel with the reduction of ROS in GCs (22). Intriguingly, a recent study suggested that the setpoint of neuronal excitability may be regulated by mitochondrial oxygen consumption capacity (67). It remains to be elucidated whether mitochondria are involved in ROS-mediated downregulation of Kv4.1 in Tg2576 mice.

## Materials and Methods

### Behavior analysis

All experiments procedures were conducted in accordance with the guide lines of University Committee on Animal Resource in Seoul National University (Approval No. SNU-090115-7). Male Tg2576 transgenic mice and their littermate wild-type control (WT) mice aged 10-weeks were trained to discriminate between two similar contexts, A and B, through repeated experience in each context. Context A (conditioning context) was a chamber (18 cm wide x 18 cm long x 30 cm high; H10-11M-TC; Coulbourn Instruments 5583, PA 18052, USA) consisting of a metal grid floor, aluminum side walls, and a clear Plexiglass front door and back wall. Context A was indirectly illuminated with a 12 W light bulb. The features of Context B (safe context) were the same as Context A, except for a unique scent (1 % acetic acid), dimmer light (50 % of A), and a sloped floor by 15° angle. Each context was cleaned with 70 % ethanol before the animals were placed. On the first 3 days (contextual fear acquisition), mice were placed in Context A for 3 min for exploring the environment, and then received a single foot shock (0.75 mA, for 2 s). Mice were returned to their home cage 1 min after the shock. On day 4 – 5, mice of each genotype were divided into two groups; one group visited Context A on Day 4 and Context B on Day 5, while the other group visited the Context B on Day 4 and Context A on Day 5. On day 4 – 5 (generalization), neither group received a shock in Context A and B, and freezing level was measured for 5 min only in Context A. We defined freezing behavior as behavioral immobility except for respiration movement. We observed video image for 2-s bouts every 10 s (18 or 30 observation bouts for 3 min or 5 min recording time) and counted the number of 2-s bouts during which the mouse displayed freezing behavior (referred to as the freezing score). The percentage of freezing was calculated by dividing the freezing score with the total number of observation bouts (18 or 30). Mice were subsequently trained to discriminate these two contexts by visiting the two contexts daily for 8 days (from day 6 to 13, discrimination task). Mice always received a foot shock (2 s) 3 min after being placed in Context A but not B. Discrimination ratios were calculated according to F_A_ / (F_A_ + F_B_), where F_A_ and F_B_ are freezing scores in Contexts A and B, respectively. To test context specificity, mice were exposed to Context A and C for 5 min each on Day 1. On Day 2, mice were placed in Context A for 3 min followed by a foot shock (2 s), and returned to their home cage 1 min after the shock. Freezing behavior was observed on the next day (Day 3) in either Context A or C for 3 min. All experiments and analyses were performed blind to the mice genotype.

### Preparation of brain slices

Brain slices were prepared from male Tg2576 transgenic mice and their littermate wild-type control (WT) mice aged from 1 to 2 months old. Average ages of WT and Tg2576 used in the present study were 5.7 week (*n* = 75) and 5.8 week (*n* = 53), respectively. Experiments for dentate GCs were mostly conducted using mice at postnatal week (PW) 4 to PW 7, while experiments for CA1-PCs were conducted using mice at PW 3 to PW 4. For *in-vivo* antioxidant pretreatment experiments, age-matched male Tg2576 mice were assigned to receive 6-Hydroxy-2,5,7,8-tetramethylchromane-2-carboxylic acid (Trolox, Sigma-Aldrich, St. Louis, MO) by intraperitoneal injection for 1 week. Trolox dosage was adopted from Betters et al. (2004), in which a 20 mg/kg priming dose was used to block oxidative stress (68). Mice were killed by decapitation after being anesthetized with isoflurane, and the whole brain was immediately removed from the skull and chilled in artificial cerebrospinal fluid (aCSF) containing (in mM): 125 NaCl, 25 NaHCO_3_, 2.5 KCl, 1.25 NaH_2_PO_4_, 2 CaCl_2_, 1 MgCl_2_, 20 glucose, 1.2 pyruvate and 0.4 Na-ascorbate, pH 7.4 when saturated with carbogen (95% O_2_ and 5% CO_2_) at 4 °C. Transverse hippocampal slices (350 μm thick) were prepared using a vibratome (VT1200S, Leica, Germany). Slices were incubated at 35 °C for 30 min and thereafter maintained at 32 °C until in situ slice patch recordings and fluorescence microscopy.

### Electrophysiological analysis for excitability and K^+^ currents

Hippocampal GCs of DG were visualized using an upright microscope equipped with differential interference contrast (DIC) optics (BX51WI, Olympus, Japan). Electrophysiological recordings were made by the whole cell clamp technique with EPC-8 amplifier (HEKA, Lambrecht, Germany). Experiments were performed at 32 ± 1 °C. The perfusion rate of bathing solution and the volume of the recording chamber for slices were 2.2 ml/min and 1.2 ml, respectively. Patch pipettes with a tip resistance of 3–4 MΩ were used. The series resistance (R_s_) after establishing whole-cell configuration was between 10 and 15 MΩ. R_s_ was monitored by applying a short (40 ms) hyperpolarization (5 mV) pulse during the recording. The pipette solution contained (in mM): 143 K-gluconate, 7 KCl, 15 HEPES, 4 MgATP, 0.3 NaGTP, 4 Na-ascorbate, and 0.1 EGTA with the pH adjusted to 7.3 with KOH. For the antibody-blocking experiments, patch pipettes were dipped into an internal solution and then back-filled with the internal solution containing the antibody of Kv4.1 at a concentration of 0.3 µg/ml. In all bath solutions (aCSF), 20 μM bicuculine and 10 μM CNQX were included to block the synaptic inputs. In voltage clamp experiments to record K^+^ currents, we added TTX (0.5 µM), CdCl_2_ (300 µM) and NiCl_2_ (500 µM) to block Na^+^ and Ca^2+^ channels, and membrane potentials were depolarized to a maximum of 30 mV for 1 s by 10 mV increments from the holding potential of −70 mV. In current clamp experiments to analyze neuronal excitability, the following parameters were measured: (1) the resting membrane potential, (2) the input resistance (R_in_, membrane potential changes (V) for given hyperpolarizing current (−35 pA, 600 ms) input), (3) AP threshold (current threshold for single action potential generation, 100 ms duration), (4) F-I curve (firing frequency (F) against the amplitude of injected currents (I), for DG; 100 pA to 600 pA, 100 pA increment, 1 s duration, for CA1 pyramidal cells (CA1 PCs); 50 pA to 250 pA, 50 pA increment, 1 s duration). Membrane potentials are given without correction for liquid junction potentials. All chemicals were obtained from Sigma (St. Louis, MO, USA), except CNQX, bicuculline, and TTX purchase from abcam Biochemicals (Cambridge, UK).

### RNA extraction and Quantitative real-time PCR (qRT-PCR)

Total RNA was isolated from mouse hippocampal DG region using TRIzol reagent (Invitrogen, Carlsbad, CA). We used 1- to 2-month-old male WT and Tg2576 mouse. cDNA was produced from 1 µg of total isolated RNA using the Invitrogen superscript III First-strand synthesis system for RT-PCR (Cat.18080-051) from Invitrogen. The expression levels of Kv4.1, Kv4.2, Kv4.3, and GAPDH were determined using sequence specific primers and Power SYBR® Green PCR Master Mix (Applied Biosystems, Part No. 4367659) for qRT-PCR, experiments were conducted on a 7900HT Fast Real-Time PCR System (Applied Biosystems). Primers used in qRT-PCR reactions as follows (5’-to −3’): GAPDH, GCAACAGGGTGGTGGACCT (forward) and GGATAGGGCCTCTCTTGCTCA (reverse), Kv4.1, GCCGCAGTACCTCAGTATCATC (forward) and GACAGAGGCAGTAGAGTTGGCA (reverse), Kv4.2, ATCGCCCATCAAGTCACAGTC (forward) and CCGACACATTGGCATTAGGAA (reverse), Kv4.3, AGATTACCACGGCCATCATC (forward) and GGAAGGAATGTTCGTGTTGG (reverse). Data were analyzed using the threshold cycle relative quantification method, with GAPDH as the endogenous control. Real-time PCR was performed in duplicate using the same amplification protocol described above.

### Western blot analysis

DG region was isolated from slices under the dissecting microscope. Isolated tissues were homogenized with a glass homogenizer in TNE buffer (50 mM Tris-HCl, pH 8.0, 150 mM NaCl, and 2 mM EDTA) supplemented with protease inhibitor cocktails (Roche), and sonicated for 10 s. After adding SDS (0.5%) and Triton X-100 (1%), lysates were incubated for 30 min at 4 °C. Insoluble materials were removed by centrifugation at 20,000 × g for 15 min at 4 °C. The amount of protein in the supernatants were determined by the BCA assay, and supernatants were mixed with 6× Laemmli sample buffer. Samples containing 20 μg protein were loaded into each lane, separated by SDS-PAGE, and transferred to a PVDF membrane. Membranes were blocked in 5 % skim milk in TBS for 1 h, and then probed with the relevant antibodies as indicated. Membranes were then incubated with peroxidase-conjugated secondary antibodies, and blots were detected with chemiluminescent reagents (Thermo Scientific). Antibodies to Kv4.1 and β-actin for Western blot analysis were purchased from Alomone lab (APC-119) and Cell signaling (#4970), respectively.

### Statistical analysis

Data were analyzed with Igor Pro (Version 6; Wavemetrics, Lake Oswego, CA) and Origin (Version 8; Microcal, Northampton, MA). All results are presented as mean ± S.E.M. with the number of cells or mice (n) used in each experiment. Statistical significance was evaluated using Student’s t-test or Mann-Whitney test, and the level of significance was indicated by the number of marks (*, P < 0.05; **, P < 0.01; ***, P < 0.001). P > 0.05 was regarded as not significantly different (N.S.). Comparison between multifactorial statistical data was made using the two-way analysis of variance (ANOVA). Differences in time-dependent changes of behavioral parameters between the two genotypes were evaluated using two-way repeated measures ANOVA.

## Abbreviations

aCSF: artificial corticospinal fluid
AD: Alzheimer’s disease
4-AP: 4-Aminopyridine
AP: Action potential
Aβ: Amyloid β
CA1: Cornu Ammonis 1 area of hippocampus
CA3: Cornu Ammonis 3 area of hippocampus
CNQX: 6-cyano-7-nitroquinoxaline-2,3-dione
DG: Dentate gyrus
fMRI: Functional magnetic resonance imaging
GABA_A_ receptors: Gamma-aminobutyric acid receptors, subtype A
GCs: Granule cells
I_A_: A-type potassium currents
I_peak_: Peak current
I_ss_: Steady state current
Kv4: Voltage-dependent potassium channel subfamily 4
L2/3: Cortical layer 2/3
LTP: Long-term potentiation
MF: Mossy fiber
qRT-PCR: Quantitative real-time polymerase chain reaction
R_in_: Input resistance
RMP: Resting membrane potential
ROS: Reactive oxygen species
shRNA: Small hairpin ribonucleic acid
TTX: Tetrodotoxin
WT: Wild-type

## Declarations

### Ethics Approval and Consent to participate

All experiments procedures were conducted in accordance with the guide lines of University Committee on Animal Resource in Seoul National University (Approval No. SNU-090115-7)

### Consent for publication

Not applicable

### Availability of data and materials

The datasets generated and/or analyzed during the current study are available from the corresponding author on reasonable request.

### Authors’ contributions

KK, YK_1_, HJ performed experiments and analyzed data. KK, JK, SL_2_, WH designed experiments. KK, WH wrote the first draft of the manuscript. SL_1_, YK_2_, WH edited the manuscript. All authors approved the final manuscript.

YK_1_: Yoonsub Kim, YK_2_: Yujin Kim, SL_1_: Sang Hun Lee, SL_2_: Suk-Ho Lee.

## Acknowledgements

Not applicable

## Authors’ information

Current address of SL_1_: Department of Neurology, Brigham & Women’s Hospital, Harvard Medical School, Boston, MA 02115, USA

